# Delayed development of basal spikelets in wheat explains their increased floret abortion and rudimentary nature

**DOI:** 10.1101/2023.02.17.528935

**Authors:** Anna Elisabeth Backhaus, Cara Griffiths, Angel Vergara-Cruces, James Simmonds, Rebecca Lee, Richard J. Morris, Cristobal Uauy

## Abstract

Wheat (*Triticum aestivum L*.) breeding efforts have increased grain yield predominantly by raising grain numbers per spikelet, rather than grain weight or spikelet number. However, across a single spike large differences exist in the number of grains per spikelet. The central spikelets produce the highest number of grains in any given genotype while apical and basal spikelets are less productive. Basal spikelets are delayed in development just after initiation and are smaller and less advanced than central spikelets already by the glume primordium stage. However, basal spikelets continue to develop and produce florets until much later in the wheat growth cycle. The precise timings or the cause of their growth cessation, and subsequent abortion, is largely unknown. In this study we investigated the underlying causes of rudimentary basal spikelet abortion. We investigated basal spikelet development in four UK winter wheat varieties as well as a set of near-isogenic lines for *VRT-A2* (*VEGETATIVE TO REPRODUCTIVE TRANSITION 2*) using shading applications in the field. We propose that basal spikelet abortion is likely the consequence of complete floret abortion as both occur at the same time and have the same response to shading treatments. Furthermore, we found that the developmental age of florets pre-abortion is an important factor for their likelihood to survive and develop viable seed. Previously, it had been proposed that reduced assimilate availability in the base of the spike leads to increased abortion. Re-analysis of published data alongside data presented here, however, does not support this model. We found that rather than assimilate availability, it is the reduced developmental age of basal florets before abortion that correlates with increased abortion. Using the floret Waddington developmental stage pre-abortion, we were able to predict final grain set per spikelet across the spike, alongside the characteristic gradient in number of grains from basal to central spikelets. We found that advancing past Waddington stage 5.5 seems to be important for floret survival and that most florets in basal spikelets had not reached this stage at the onset of floret abortion. The abortion of all florets could therefore be the reason for their rudimentary appearance in the mature spike, suggesting that basal spikelet abortion is simply the consequence of all florets inside the spikelet being aborted and thus all other spikelet structures (e.g., lemma, rachilla, glume) also ceasing to develop. Future efforts to improve spikelet homogeneity across the spike could thus focus on improving basal spikelet establishment and increasing floret development rates pre-abortion.

## Introduction

Three of the globally most important staple crops, maize (*Zea mays L*.), rice (*Oryza sativa L*.) and wheat (*Triticum aestivum L*.), belong to the family of grasses (*Poaceae*). One characteristic feature of the *Poaceae* is their inflorescence, which develops florets inside specialised structures termed spikelets (Kellogg, 2001; Kellogg et al., 2013). Upon floral transition the apical meristem of grasses elongates and, depending on the species, the spikelets are formed directly on the rachis (e.g., wheat) or on primary (e.g., maize) or secondary (e.g., rice) branches (Kellogg, 2022). Each spikelet can form multiple florets, however in some species the number of florets per spikelet is highly genetically controlled (e.g., maize) while in others the number is also dependent on environmental factors (e.g., wheat) (Bonnett, 1966). Across all grasses the potential yield of a plant depends on the number of inflorescences, number of spikelets and florets per inflorescence, as well as grain weight.

Over the past century, breeding efforts have enhanced yields in winter wheat, predominantly through an increase in grain number, rather than grain weight (Würschum et al., 2018; Voss-Fels et al., 2019; Sakuma and Schnurbusch, 2020). Furthermore, it has been established that the increase in grain numbers was achieved through the improvement of grains per spikelet rather than through an increase in spikelets per inflorescence (termed spike in wheat)(Philipp et al., 2018; Sakuma and Schnurbusch, 2020). However, increasing the number of grains per spike can have negative effects on grain weight, as is the case of ‘Miracle Wheat’ (Poursarebani et al., 2015). Previous research found that the amount of resource available to the plant can affect the relative growth and development of the initiated spikelets and grains. Thus, trade-offs between the different yield components are created by the ‘source-sink’ balance (Reynolds et al., 2009).

During wheat spike development, a finite number of spikelets is initiated until terminal spikelet formation, thus the number of spikelets per spike is determined relatively early in the crop growth cycle (Bonnett, 1966; Kirby and Appleyard, 1981). Each spikelet initiates an indetermined number of floret primordia, of which most are aborted. Wheat spikelets initiate typically many florets (10-12), but only a fraction (typically 3-5) survive abortion and go on to form grains (Sadras and Slafer, 2012). Thus, a higher number of spikelets per spike might not lead to more grains per spike as it can be annulled by increased floret abortion. Furthermore, the weight of individual grains has also been shown to correlate negatively with the number of grains (Sakuma and Schnurbusch, 2020).

Over the past decade, several genes that affect spikelet number, floret abortion and grain weight have been identified in wheat. For example, several genes that increase the number of spikelets initiated have been cloned, such as *FRIZZY PANICLE (FZP)* (Dobrovolskaya et al., 2015; Poursarebani et al., 2015)*, WHEAT ORTHOLOG OF APO1 (WAPO-A1)* (Kuzay et al., 2019; Muqaddasi et al., 2019; Voss-Fels et al., 2019), and *TEOSINTE BRANCHED1 (TB1)* (Dixon et al., 2018). However, the introgression of the beneficial alleles into elite material has seldom led to significant increases in yields. For example, the increase in expression of *WAPO1* leads to increased spikelet numbers, which does not translate into yield gains due to increased spikelet abortion (Wittern et al., 2022) and decreased floret survival (Kuzay et al., 2022). Targeting the number of florets rather than spikelets, the reduced-function allele of *GRAIN NUMBER INCREASE 1* (*GNI1*) has been shown to confer yield increases by reducing floret abortion compared to the wildtype *GNI1* allele, which functions as a rachilla growth inhibitor (Sakuma et al., 2019). The high frequency (96%) of the increased grain number *GNI1* allele among durum wheats suggests that there has been a strong selection pressure for increased grains per spikelet during domestication and breeding (Sakuma et al., 2019). In terms of grain weight, ectopic expression of a semi-dominant allele of the *VEGETATIVE TO REPRODUCTIVE TRANSITION 2* (*VRT-A2*) gene from *T. turgidum* ssp. *polonicum* leads to increased grain weight, yet does not affect grain number per spike nor increase yield with respect to the wildtype *VRT-A2a* allele across multi-year field trials (Adamski et al., 2021).

Within a spike, large differences in the number of grains per spikelet (i.e., spikelet fertility) exist, with the central spikelets producing the highest number of grains in any given genotype. An analysis of grain distributions across the spikes of 210 elite and 180 heritage wheat accessions showed that the number of grains per spikelet has increased in elite material, but that this increase has mostly occurred in the central spikelets, to a lesser extent in apical, and not at all in the most basal spikelets (Philipp et al., 2018). The authors proposed that reducing the variation in spikelet fertility across the spike could be a promising avenue to increase yields and improve grain size homogeneity. However, we have little understanding of the factors that determine spikelet productivity gradients within a spike.

Apical spikelets are initiated last and thus have less time to develop their floret primordia (Bonnett, 1966). The most basal spikelets are initiated first, but they often develop only in a rudimentary form, meaning that they are much smaller than other spikelets, produce no grain and have underdeveloped glumes and lemma. This variation in spikelet development leads to the characteristic lanceolate shape of the wheat spike (Backhaus et al., 2022). Basal spikelets are delayed in development just after initiation and are smaller and less advanced than central spikelets already by the glume primordium stage (Bonnett, 1966). However, the basal spikelets continue to develop and produce florets until later in spike development when basal spikelet abortion occurs. The precise timings or the cause of their growth cessation, and subsequent abortion, is largely unknown in wheat.

A wealth of experimental data has confirmed that across all spikelets resource availability, also termed source strength, is closely linked to floret survival (González et al., 2011; Ferrante et al., 2013). As a way to explain the higher floret abortion of basal spikelets, González et al. (2011) proposed that basal spikelets have poorer resource allocation than central spikelets, although it remains to be established whether basal spikelets do indeed have lower priority in assimilate partitioning than central spikelets. Other factors, such as development stage of florets (Ferrante et al., 2020), vasculature development (Hanif and Langer, 1972) and distance to the rachis (Kadkol and Halloran, 1988) have also been shown to affect floret survival in spikelets, but none of these factors have been investigated further as the cause of rudimentary basal spikelets.

Multiple studies have used shading applications to reduce photosynthetic activity in the field and shown that pre-anthesis shading reduces yield and also spike and plant dry weight, which is a good indicator of reduced source strength (Fischer, 1985; Savin and Slafer, 1991; Slafer et al., 1994). The effects of shading on altering resource availability are relatively quick (within two days) as determined by measurements of water-soluble carbohydrates (WSC; Stockman et al., 1983). Stockman et al. (1983) also found that shading treatments affected basal spikelet fertility more than apical and central spikelets (using single tiller plants under controlled environment conditions). Furthermore, Slafer et al. (1994) reported the effect of shading on each individual spikelet and showed that the number of basal spikelets with zero fertile florets was increased by 3-4 spikelets under shading conditions. This suggests that not only floret, but also basal spikelet survival, is negatively affected by shading conditions.

In this study, we aimed to characterise the causes of rudimentary basal spikelet development in wheat. We used shading treatments pre-anthesis to reduce resource availability in precise and short time frames that spanned basal spikelet abortion during the crop cycle. We collected samples after the shading application to assess the effect of shading on sugar concentrations in different spikelet positions across the spike. We also traced the development and number of florets across different spikelet positions to relate pre-anthesis floret development to the probability of floret survival. This study highlights that rudimentary basal spikelets are most likely a consequence of complete floret abortion in basal spikelets. We did not find any evidence for lower assimilate accumulation in the base, but rather that the delayed development of the florets in basal spikelet can explain to a large extent their abortion and rudimentary nature.

## Results

### Basal spikelet development ceases two weeks pre-anthesis and is sensitive to resource availability at that time

To investigate when basal spikelet abortion takes place, we applied shading treatments that reduce assimilate availability in field grown wheat plots at defined growth stages (Kemp and Whingwiri, 1980; Stockman et al., 1983). Each shading treatment consisted of ca. 45% light reduction over 12/13 days in field grown plots (Supplemental Table S1). In 2021, we applied two shading treatments; the first treatment (Shading A) started around the stem extension phase, whereas the second treatment (Shading B) was applied one day after removal of Shading A and ended ca. 10 days before anthesis (Figure 1A&B). We applied shading to four UK winter wheat cultivars as well as a set of cv. Paragon near isogenic lines (NILs) carrying either the wildtype *VRT-A2a* or the *T*. *polonicum VRT-A2b* allele (Adamski et al., 2021). *VRT-A2b* has been previously shown to increase the number of rudimentary basal spikelets (RBS) by 1-2 spikelets compared to the Paragon *VRT-A2a* sibling NIL (Backhaus et al., 2022). We found that the early timeframe in 2021 (Shading A) had no effect on the number of RBS formed, however Shading B increased the number of RBS significantly across all genotypes by on average 1.46 RBS (Figure 1C&D). As both shading treatments were applied for the same duration, we hypothesised that basal spikelet abortion is more sensitive to source alterations between 10 and 22 days before anthesis.

**Figure 1:**
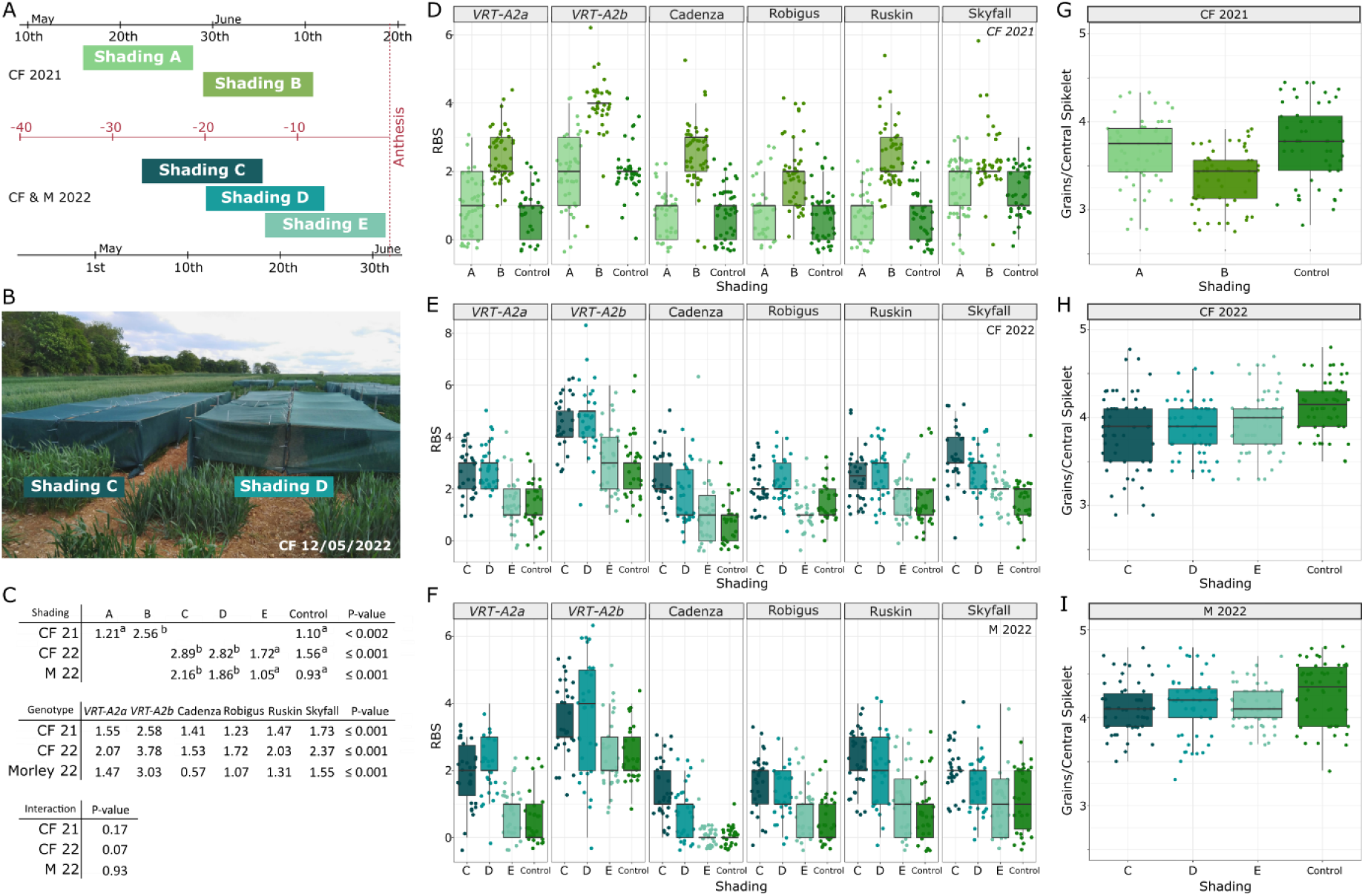
Effect of the different pre-anthesis shading treatments on spike traits. (A) Timing of shading applications in Church Farm (CF) in 2021 and CF and Morley (M) in 2022 relative to the average date of anthesis (Supplemental Table S2). Shading A and B were applied for 13 days each (2021), shading in 2022 was applied for 12 days. (B) Image of shading nets for Shading C and D in the field. (C) Estimated means of rudimentary basal spikelet (RBS) for the different shading treatments (top) and genotypes (middle). ANOVA test of significant difference was performed for each trial independently and letters (a-b) indicate LSD results. Interaction between genotype and treatment was non-significant (NS) in CF 2021 and Morley 2022, and borderline NS in CF 2022 (bottom). (D-F) Number of RBS per spike recorded for ten spikes from each block (N=3) in control vs shading applications in CF 2021 (D), CF 2022 (E) and Morley 2022 (F). (G-I) Number of grains per spikelet in the central most spikelets (spikelet position 10-12) from the same data trials. Box represents the middle 50% of data with the borders of the box representing the 25th and 75th percentile. The horizontal line in the middle of the box represents the median. Whiskers represent the minimum and maximum values, unless a point exceeds 1.5 times the interquartile range in which case the whisker represents this value. Points in D-F represent all subsamples (individual RBS measurements from ten to twenty spikes of each of the three blocks), whereas statistical analyses were performed with mean values. In G-I points represent the average number of grains/spikelet of the central three spikelets taken from ten individual spikes from each block. (Raw data can be found in Supplemental Dataset S1).

To expand on these results and further investigate when RBS are determined, we replicated the experiment in 2022 across two locations (Church Farm (CF) and Morley (M)). In both locations we applied three shading treatments, which overlapped by one week, allowing us to further narrow down the timing of rudimentary basal spikelet formation (Figure 1A). The earliest shading treatment (Shading C) was applied during the stem extension phase and the last shading treatment (Shading E) finished at anthesis. Across all genotypes, Shading C and Shading D significantly increased RBS numbers by 1.33 and 1.26 RBS in CF, respectively, and by 1.23 and 0.93 RBS in Morley, respectively (Figure 1E-F). Shading E had no significant effect in CF (Figure 1E) and Morley (Figure 1F). This suggests that the last week of Shading C, which was also the first week of Shading D, overlapped the timeframe in which RBS formation is most sensitive to resource limitations, in this case between 10 and 16 days pre anthesis. Shading E on the other hand was applied after basal spikelet abortion had happened and the number of RBS had been determined. This critical timeframe in 2022 is consistent with the 2021 results and supports the idea that rudimentary basal spikelet formation is linked to a specific growth stage ca. two weeks pre-anthesis.

In addition, we recorded mature plant weight in both Church Farm trials and the number of spikelets/spike in all three trials. In 2021, both shading applications significantly reduced plant weight compared to the control (Supplemental Table S3). In 2022, all shading applications reduced plant weight, although not significantly (Supplemental Table S3). The reduced final plant weight confirms the expected effect of shading on resource availability. Furthermore, the effect of shading on plant weight was equivalent for all shading treatments within the trial and thus any differences in shading effects on RBS between the treatments would be due to timing, rather than intensity. The number of spikelets per spike was not affected by our shading treatments (Supplemental Table S3), which is consistent with the fact that the treatments were applied post the terminal spikelet phase and spikelet initiation was already completed (Kirby and Appleyard, 1981). This also eliminates the possibility that RBS numbers increased due to more spikelets being initiated.

Across the six genotypes, the effect of shading was consistent, only a borderline non-significant interaction (*P* = 0.07) between genotype and shading was detected in CF 2022 were Shading E also slightly increased RBS in cv. Ruskin (Figure 1E, Supplemental Table S3). Consistent with our previous results, the introgression of *VRT-A2b* increased the number of RBS by 1-2 in all three trials and showed a linear response under shading conditions (Figure 1C, Supplemental Table S3). Furthermore, some of the significant differences in number of RBS between the winter wheat varieties were consistent across the three trials. For example, across all trials cv. Skyfall had a significantly higher number of RBS (1.6-2.4) than cv. Cadenza (0.6-1.5), which overall had the lowest number of RBS together with cv. Robigus (Figure 1C). This suggests that genetic components beyond *VRT-A2* affect RBS in UK wheat varieties and that these could be further studied in the future.

To investigate if the critical timeframe for rudimentary basal spikelet formation coincides with the well-defined timeframe of floret abortion, we recorded the number of grains per spikelet in the central and most fertile spikelets of the spike. In 2021, the number of grains per central spikelet was significantly reduced by Shading B (*P* = 0.007), but not by Shading A. In 2022, the number of grains per spikelet were reduced less, but significantly by shading C and D in CF (*P* = 0.05), but not in Morley (*P* = 0.28). Across all three trials, the central spikelet fertility was not significantly different between *VRT-A2* NILs, suggesting that floret abortion is not increased by the *VRT-A2b* allele. The results from 2021, and to a lesser degree from 2022, suggest that basal spikelet abortion might be happening at the same time as floret abortion in the central spikelets and we hypothesised that both are possibly controlled by the same mechanisms.

### Complete floret abortion likely causes rudimentary basal spikelet formation

To investigate the hypothesis that basal spikelet and floret abortion are determined during the same development phase, we harvested and dissected spikes during shading treatments in both 2022 trials (Morley and CF) and recorded floret number and Waddington development stage in the basal six and central two spikelets. In the control conditions at both locations, the number of florets per spikelet increased in all genotypes from 27 to 20 days pre anthesis, except in Skyfall, which had a similar number of living floret primordia between these two stages. At day 20 pre-anthesis, we recorded the maximum number of florets across the time course, varying between 9-10 (interquartile range), with a maximum of 11 florets/spikelet across all genotypes and both locations (only control plots analysed). Previous studies reported similar numbers of maximum florets in winter wheat (Guo et al., 2016), which suggests that in this experiment the second sampling timepoint (20 days pre-anthesis) overlapped the maximum floret stage of wheat spike development. In the following week, the number of living floret primordia per spikelet decreased drastically by 4-6 (interquartile range) florets per spikelet (Supplementary Table S4). From 13 to 0 days pre-anthesis, the number of living floret primordia per spikelet decreased only slightly. Our results align with the findings of previous studies that described the pattern of floret initiation and abortion over the wheat growth cycle (González et al., 2011; Guo et al., 2016).

Shading C and Shading D both overlapped the critical timeframe of floret abortion (20-13 days pre-anthesis) while Shading E was applied post floret abortion phase. This strengthens our hypothesis that shading only affected RBS numbers when applied during the floret abortion phase (Figure 1 D-F). If shading is applied before (Shading A, 2021) or after (Shading E, 2022) the floret abortion phase, the impact of shading on basal spikelet abortion was not significant.

Comparing the number of living floret primordia between basal and central spikelets, we observed that onset of abortion is synchronised across all six basal and the two central spikelets around 20 days pre-anthesis. The number of living floret primordia at/before abortion is relatively similar across all spikelets (overall mean = 9.13; Supplemental Table S4), although the number of living floret primordia is highest in the 2 central and upper 3 basal spikelets (mean= 9.57) and decreases gradually, albeit significantly, from the third basal spikelet (8.92) to the second (8.5) and most basal spikelet (7.79; Supplemental Table S4). Furthermore, floret abortion seems to be more intense in the most basal 2 spikelets, which lose proportionally more florets during the abortion phase than all other spikelets. While central (C1-C2) and upper basal spikelets (B3-B6) lose on average 5.33 florets (or 56% of the initiated florets), the most basal two spikelets abort significantly more florets, 6.88 out of the 8.5 initiated (81%) in the second most, and 7.12 of the 7.79 initiated florets (91%) in the most basal spikelet (Supplemental Table S4). Thus, it is the lower initiation of living floret primordia pre-abortion and the increased loss of florets during abortion that leads to the loss of all florets in basal spikelets in several of the genotypes. The abortion of all florets could be the reason for their rudimentary appearance in the mature spike, suggesting that basal spikelet abortion is simply the consequence of all florets inside the spikelet being aborted and thus all other spikelet structures (e.g., lemma, rachilla, glume) also ceasing to develop any further. This would lead to their small and underdeveloped, rudimentary appearance at maturity.

It remains unclear, however, why floret abortion affects basal spikelets more severely. Previous studies hypothesised that basal spikelets would have less priority in assimilate partitioning than the central and apical spikelets (Stockman et al., 1983; González et al., 2011). To test this hypothesis, we measured the concentration of sugars across the spikes collected from the control and shading plots at the end of Shading B (2021) and Shading D (2022). We dissected the spikes into basal and central sections for both spikelets and rachis to investigate differences in sugar concentrations across the spike. As spikelets are already varying in size and development at this point, we also analysed sugar concentrations in the rachis, which is more stable in size across the spike. In general, spikelets had significantly higher sugar concentrations compared to rachis sections (Supplemental Table S5). When comparing between the basal, central and apical sections, we generally found no significant differences in sugar concentrations (Table 1, Supplemental Table S5), albeit we did detect cases of significant differences between sections for all three sugars (fructose, glucose and sucrose) in specific treatment/tissue combinations. However, in *all* these cases the basal section had higher concentrations compared to the central section (see Supplemental Table S5, for example sucrose in M2022). We thus found no evidence for basal tissues (spikelets or rachis), having lower sugar concentrations than the central tissues.

**Table 1:** Sugar concentrations (μg/mg tissue weight) after shading treatments across the spike. Values are upper (UCI) and lower (LCI) confidence interval, the mean of the data is thus the mid-point between the two values. Raw data can be found in Supplemental Dataset S3. All statistical analyses are detailed in Supplemental Table S5.

|  |  |  | CF 2021 |  | M 2022 |  |  | CF 2022 |  |  |
| --- | --- | --- | --- | --- | --- | --- | --- | --- | --- | --- |
|  |  |  | Glucose | Fructose | Glucose | Fructose | Sucrose | Glucose | Fructose | Sucrose |
| Control | Spikelet | Apical | - | - | 26.4-32.8 | 17.9-21.6 | 17.9-21.2 | 19.4-24.8 | 13.1-17.1 | 13.0-18.1 |
|  |  | Central | 12.4-13.8 | 8.3-10.4 | 24.5-27.4 | 18.7-20.6 | 18.0-20.3 | 18.4-29.4 | 12.0-20.6 | 10.8-19.6 |
|  |  | Basal | 11.2-13.8 | 8.0-9.8 | 22.9-31.6 | 16.7-24.4 | 16.0-23.4 | 17.1-21.4 | 11.4-15.4 | 10.2-16.3 |
|  | Rachis | Apical | - | - | 15.3-26.3 | 12.3-19.0 | 13.8-19.5 | 11.7-19.0 | 9.6-16.1 | 10.3-17.6 |
|  |  | Central | 8.5-10.2 | 6.7-10.3 | 16.8-18.3 | 14.1-15.6 | 14.7-17.0 | 13.2-16.7 | 10.8-14.3 | 11.5-15.6 |
|  |  | Basal | 10.4-13.3 | 7.7-9.7 | 18.9-24.2 | 15.2-18.0 | 16.7-19.2 | 15.5-18.3 | 11.2-13.6 | 10.6-15.8 |
| Shading | Spikelet | Apical | - | - | 24.5-27.5 | 16.5-19.1 | 15.6-17.3 | 19.6-26.4 | 13.0-17.9 | 12.5-17.2 |
|  |  | Central | 10.7-12.7 | 5.8-8.8 | 24.7-26.4 | 18.4-19.3 | 16.7-18.1 | 22.9-27.5 | 16.2-19.9 | 15.6-18.8 |
|  |  | Basal | 9.6-12.8 | 5.4-8.2 | 22.4-32.8 | 16.2-23.2 | 14.7-21.3 | 21.5-27.7 | 14.4-19.0 | 13.7-17.6 |
|  | Rachis | Apical | - | - | 12.2-20.1 | 8.5-14.3 | 8.2-13.8 | 15.4-26.5 | 10.9-18.6 | 11.1-17.7 |
|  |  | Central | 7.0-8.6 | 3.6-5.8 | 16.1-19.5 | 12.5-14.9 | 12.5-14.7 | 12.3-18.4 | 8.8-14.5 | 9.4-15.0 |
|  |  | Basal | 8.8-11.6 | 3.7-5.6 | 21.1-26.1 | 15.4-18.3 | 14.9-17.8 | 20.0-23.3 | 14.5-16.6 | 15.2-16.9 |

### Florets in basal spikelets are less developed than same florets in central spikelets

Rather than only considering total living floret primordia per spikelet at 20 days pre-anthesis, we wanted to investigate how developmental age of each floret at this timepoint affects their survival chance. To address this, we compared the development stages of each floret from the most basal floret (F1) to the most distal floret from the rachis (F8) across the basal six and central two spikelets (Figure 3A; Supplemental Table S6). By taking the mean developmental age of the florets from all genotypes in the CF trial pre-abortion (20 days pre-anthesis), we found that florets in basal spikelets are less developed than their central spikelet counterparts (Figure 3A; Supplemental Table S6). For example, floret F1 in the basal spikelets had on average reached Waddington stage 5.4 at this timepoint while the same floret in the central spikelets had on average reached Waddington stage 6.5 (Figure 3A; Supplemental Table S6). This additional information is not available when recording only the number of living floret primordia per spikelet as has been the case in previous studies (Stockman et al., 1983; Sibony and Pinthus, 1988; Craufurd and Cartwright, 1989). Comparing the development of the equivalent floret positions (e.g., F1 to F1, F2 to F2) across the spike reveals that all florets in the basal spikelets (B1) lag behind their central counterparts (C2) by approximately one Waddington stage (1.02 ± 0.04 SE).

**Figure 2:**
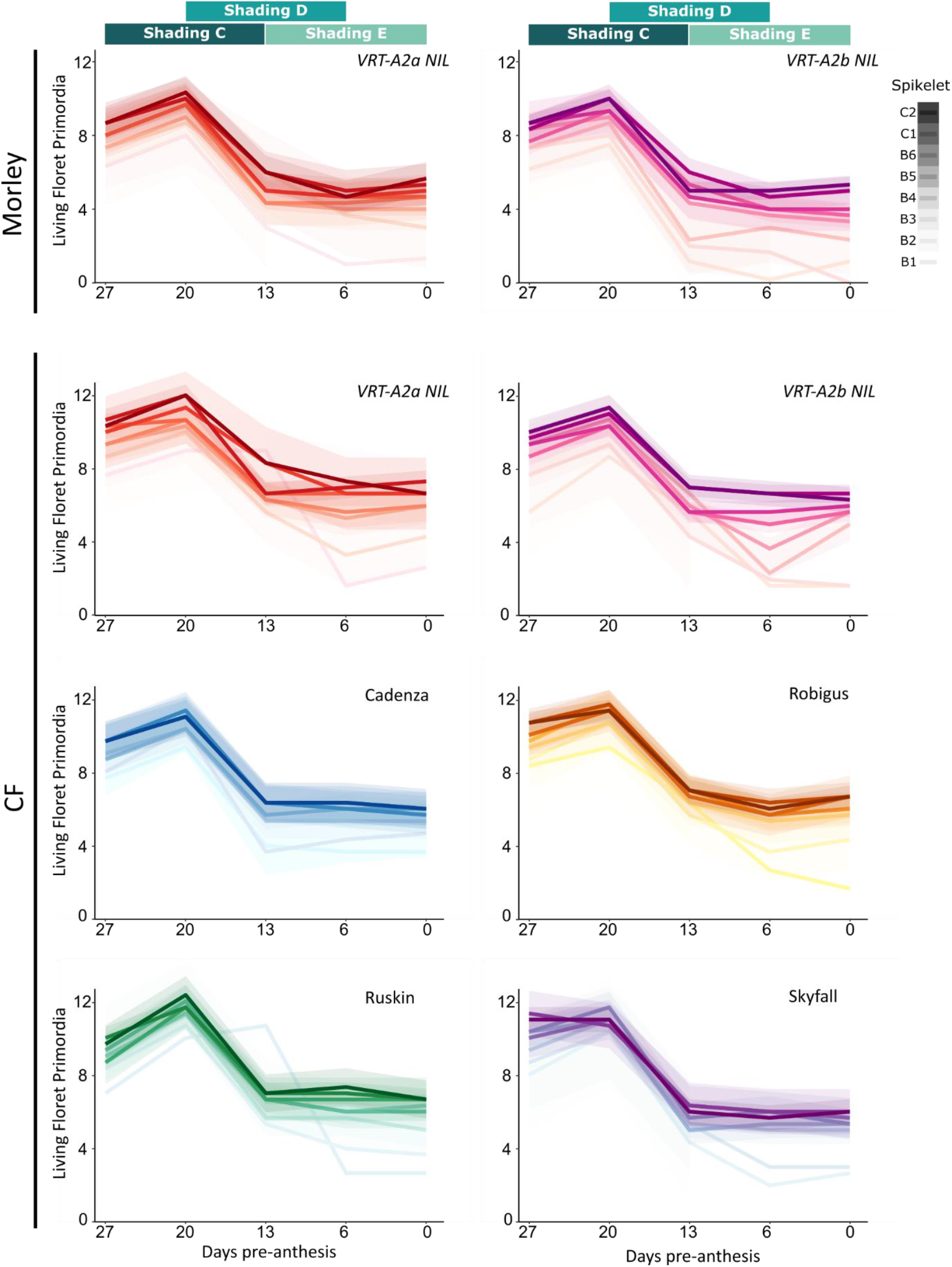
Floret fertility pre-anthesis in control conditions. We counted the number of living floret primordia per spikelet (y-axis) once per week from 27 to 0 days before anthesis (x-axis), with the first sampling coinciding with the beginning of Shading C. Each week we collected spikes from the control condition (1 spike per block, n=3) and dissected the six basal (B1-B6) as well as the two central (C1-C2) spikelets. In CF 2022, we analysed all genotypes, but in Morley 2022 only the *VRT-A2* NILs were sampled. Green boxes on top of the graphs represent the timing of the three overlapping shading applications. Colour intensity of the line indicates the spikelet position along the spike (darkest = most central spikelet). Shaded areas represent 95% confidence intervals for each spikelet. (Raw data can be found in Supplemental Dataset S2).

**Figure 3:**
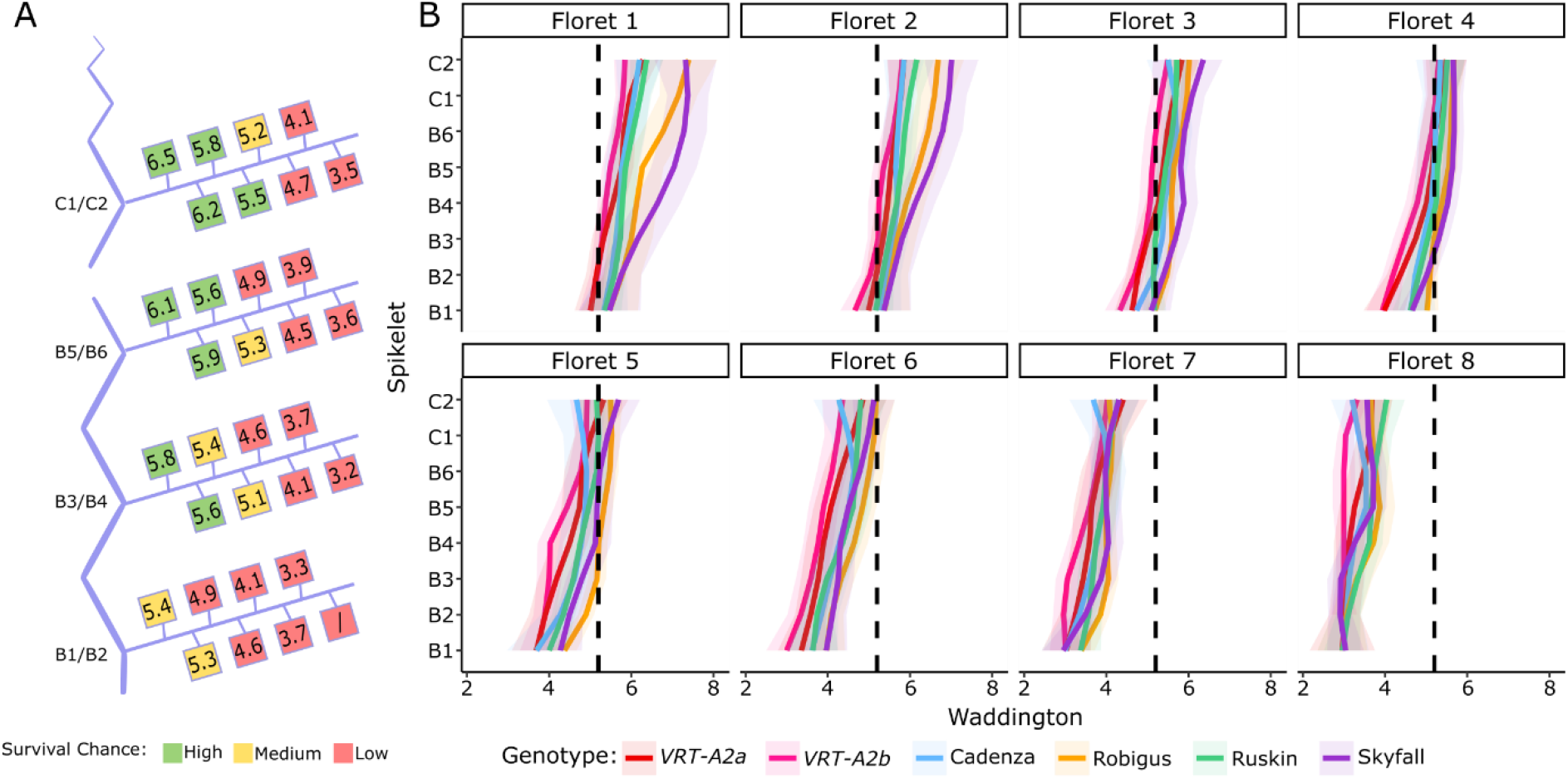
Waddington development stage of florets across the spikes 20 days pre-anthesis. (A) Graphical representation of the spike with the basal (B1-B6) and central (C1-C2) spikelets analysed in this study branching off the main rachis (vertical blue). Within each spikelet position, the average developmental score of two distichous spikelets across all genotypes (in CF 2022, Control) is represented by the coloured squares from the most proximal floret (F1, left) to the most distal floret (F8, right; Supplemental Table S6). Survival chance estimation is indicating by colouring, florets with Waddington stage ≥ 5.5 are green, between W5 and W5.5 yellow and below W5 are red. (B) Waddington stage of florets F1 to F8 across the spike, from basal (B1-B6) to central (C1-C2) spikelets. Colours represent genotypes. The black dotted line is positioned at Waddington stage 5.25 (as an arbitrary transition point of survival chance). Shaded area represents 95% confidence interval.

It is generally accepted that floret survival after abortion, rather than floret number pre-abortion, is the determining factor of final grains per spikelet (Langer and Hanif, 1973; Fischer and Stockman, 1980; Whingwiri and Stern, 1982; Sibony and Pinthus, 1988; González et al., 2003, 2005; Ferrante et al., 2010). These studies, however, did not consider differences in development pre-abortion. We therefore wanted to test if the differences in developmental age of the basal spikelets might be responsible for the greater floret abortion and subsequently, lower grain numbers. To explore this idea, we hypothesised that; (1) the more developed florets are, the less likely they are to be aborted and that, (2) florets therefore need to have reached a minimum Waddington stage to have a high chance of survival. Using data collected at maturity (final grain number per spikelet; Supplemental Table S7), we determined that under field conditions in the 2022 trials, florets beyond Waddington 5.5 at 20 days pre-anthesis had a very high chance of survival whereas florets below Waddington 5.0 had a low chance of survival. To illustrate this concept, we coloured the florets in Figure 3A as either red (low survival chance, Waddington < 5.0), yellow (medium survival chance, 5.0 ≤ Waddington < 5.5) or green (high survival chance, Waddington ≥ 5.5). Using this criterion, we would predict that across all spikelet positions, florets F6, F7 and F8 have a very low survival chance as they do not pass this threshold, while floret F5 would have a medium chance of survival only in central spikelets and in some genotypes (Figure 3A and 3B). The most basal florets (F1 and F2) would be assumed to have a very high chance of survival in all spikelets except the most basal spikelets (B1 and B2). It is important to note that this the threshold of survival at Waddington 5.5 was chosen as it best fits the actual number of grains per spikelet observed in mature spikes (ca. 4 grains per central spikelet, Figure 1H-I).

### Differences in development pre-abortion can be used to predict reduced grain numbers in basal spikelets

The apparent relationship between florets above W5.5 and final grain number raises the question whether the developmental age of florets at the onset of abortion affects their likelihood to survive the abortion process itself. To test this hypothesis, we predicted the number of grains per spikelet for each genotype in control conditions, using the floret data pre-abortion (20 days pre-anthesis). Based on its Waddington stage, we assigned a probability of survival (a cumulative normal distribution function, CDF, with a mean = 5.5, sd = 0.195) for each floret to avoid abortion and produce grain. This allowed us to assign a ‘survival probability’ to each floret regardless of its position within the spike or spikelet. The survival probability increases as development advances and this is captured well by a cumulative normal distribution (or similar saturation functions such as the Hill or logistic function). For example, the survival probability of florets in Waddington stage 5.5 is 0.5, while the survival probability for florets in Waddington stage 6 is nearly 1 (0.99) and for florets in stage 4.5 it is close to 0 (1.4 x 10^-7^). Summing over these probabilities, rather than counting the number of florets above W5.5, allowed us to compute the expected number of florets that survive whilst accounting for a degree of uncertainty in the survival rate of florets based on the Waddington score. Using this method, we calculated the floret survival probability for florets 1 to 8 within a spikelet and then summed up the probabilities, leading to a predicted grain number per spikelet (Figure 4A, light grey). Comparing these values to the actual number of grains per spikelet recorded in mature spikes (Figure 4A, solid colours) reveals a close fit. This approach predicted successfully for basal spikelets to have the least grains/spikelet, and for grain numbers to increase steadily towards the central spikelets, as is the case in the mature spike data. The correlation between the actual and predicted grain numbers was high (0.81) and the y-intercept was close to 0 (Figure 4C). Using only the number of florets per spikelet pre-abortion, without considering their Waddington stages, fails to predict the gradient across the spike and has consequently a much worse fit to the actual data (Supplemental Figure S1).

**Figure 4:**
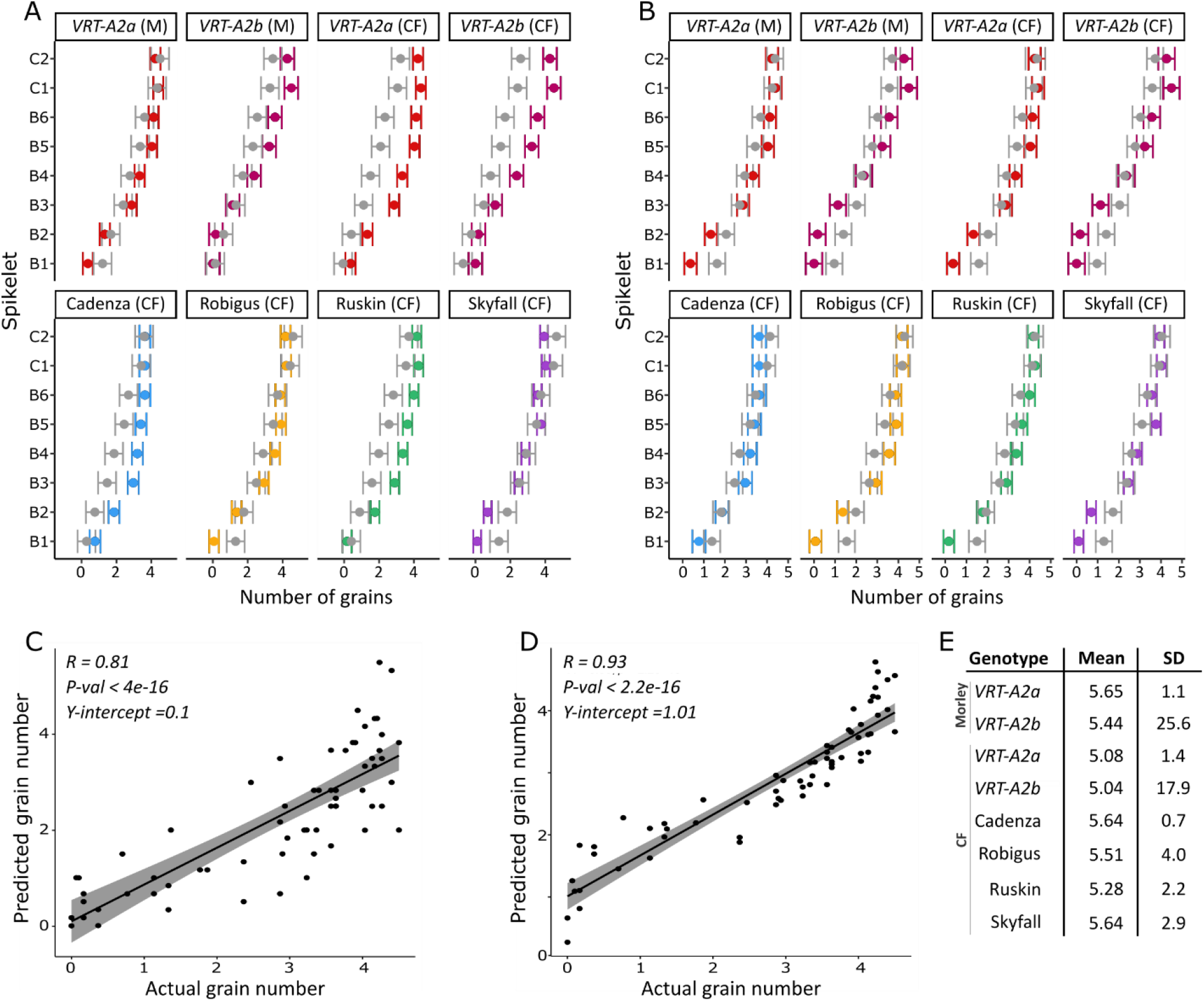
Prediction of grains per spikelet from floret development pre-abortion. (A) Number of grains per spikelet recorded in mature spike (dark colours) and predicted number of grains, resulting from the expected number of survived florets for florets 1-8, calculated from a cumulative normal distribution with a half-maximal value at the Waddington stage of 5.5 (mean) and a survival decay characterised by a standard deviation of SD = 0.195 (grey). (B) Number of grains predicted per spikelet*Genotype after optimisation of the mean and standard deviation of the normal distribution for each genotype individually (grey) versus actual grain numbers recorded at maturity (colour). Data in A and B is shown confidence interval (bars) plus mean (dot). (C) Linear regression and fit of Actual grain number versus predicted grain number using data from (A). (D) Linear regression and fit of actual versus predicted grain numbers using data predictions shown in (B). (E) Resulting mean and SD values for each Genotype*Experiment from function optimisation by genotype (Data in panel A&D). B = Basal (spikelet), C = Central (Spikelet). Raw Data for actual grains/spikelet can be found in Supplemental Dataset S1, predicted number of grains/spikelet can be found in Supplemental dataset S4).

Using a t-test, we found that there are no significant differences between the actual and predicted grain values for most of the genotypes. The predictions that fit least well were for Cadenza (*P = 0.10*) and *VRT-A2b* (*P = 0.10*) (Supplemental Table S7). Similarly, not all spikelets were fitted equally well, and for the 4^th^, 5^th^ and 6^th^ spikelet from the base, predictions were significantly lower from the actual grain values (Supplemental Table S7, *P < 0.05*). Next, we explored if the mean and SD of the CDF could be optimised individually for each genotype to achieve a better fit of the actual data. For this we again used the cumulative normal distribution probability function but optimised the mean and SD fully unconstrained using simulated annealing and the downhill simplex method (both gave consistent results). The resulting predictions significantly matched the actual number of grains/spikelet (Figure 4B) and improved the overall correlation to R = 0.93 (Figure 4D). When comparing the optimised predicted grain values to the actual grain values per genotype, we found no significant nor borderline significant differences. The optimisation led to each genotype having an individual mean (Waddington) stage as turning point for survival probability (Figure 4E).

Despite achieving an overall better fit, the correlation between actual and optimised predicted grain values had a higher y-intercept (1.01), which suggests that the approach slightly over-predict low grain values. This was also confirmed by the t-test, which found significant differences between the actual and predicted values for the 1^st^ and 2^nd^ most basal spikelet (Supplemental Table S7), especially in Robigus and Ruskin. Overprediction of the lower grain values might be due to the optimisation algorithm favouring a better fit of central and upper spikelet values as the errors are larger in proportion and because we included more spikelets with higher grain values, leading to an unequal distribution in datapoints (Figure 4C&D). More data over the full range will allow us to test this hypothesis and validate the proposed model.

## Discussion

### Basal spikelet abortion is likely the consequence of complete floret abortion

In wheat and other grasses, the most basal spikelets are generally less productive than central spikelets and are often only developed in a rudimentary form. Basal spikelets are initiated first, yet the number of grains per spikelet is lowest in the most basal spikelets and increases gradually towards the centre of the spike. Although previous studies have highlighted that basal spikelets are less productive and more readily aborted under stress conditions (Stockman et al., 1983; Savin and Slafer, 1991; Ferrante et al., 2020), the causes of rudimentary basal spikelet formation have not been studied in detail. Previously, we found that basal spikelets are delayed in growth and development immediately after their initiation due to lower expression of spikelet initiation genes than in central spikelets (Backhaus et al., 2022). However, basal spikelets continue to grow and develop throughout the crop cycle and their rudimentary status is finalised later. In this study, we investigated the timing and physiological mechanisms of basal spikelet cessation.

We found that applying resource-limiting shading treatments in the field between 13 to 20 days pre-anthesis significantly increased rudimentary basal spikelet (RBS) numbers by between 0.93 to 1.46 in three independent experiments (Figure 1). We therefore hypothesised that basal spikelet abortion is happening during this very defined timeframe and is highly sensitive to resource availability in this period. The timeframe identified for basal spikelet abortion overlaps stem elongation phase and has previously been termed the ‘critical period’ of wheat development (Fischer, 1985; Siddique et al., 1989; Savin and Slafer, 1991). During the critical period, the stem and spike are undergoing maximal growth and are rapidly accumulating biomass, which has been proposed to be a competitive process between the stem and spike (Fischer and Stockman, 1980; Siddique et al., 1989). Furthermore, floret abortion also takes place during the 10-20 days pre-anthesis. Previous studies showed that shading during the critical period significantly decreases the number of grains per spikelet (Stockman et al., 1983; Savin and Slafer, 1991; Slafer et al., 1994). Thus, through a variety of studies in which floret fertility was manipulated using genetic or environmental factors, the initiation of floret abortion has been tightly linked to the stem extension phase where resource availability is directly, or indirectly, determining the number of florets that will survive abortion. In 2021, we found that shading B significantly reduced central spikelet fertility and in the 2022 experiments, shading C and D had a negative, although not significant, effect on central spikelet fertility (Figure 1G-I). Dissections of the spike during the shading treatments in 2022 confirmed that the two shading treatments leading to increased number of RBS (C and D) overlapped the week of floret abortion, while shading E was applied post the floret abortion phase (Figure 2). We therefore concluded that shading increases basal spikelet abortion only if applied during the ‘critical period’ of spike growth and floret abortion.

As basal spikelet abortion is happening at the same time as floret abortion and is affected similarly to floret abortion by shading, it raises the question if spikelet abortion is simply the consequence of all florets being aborted in these spikelets. The nearly complete abortion of florets in the most basal spikelet across all genotypes supports this idea. Unlike spikelets, the number of florets is indeterminate and each spikelet initiates many floret primordia (in this study on average 9 florets/spikelet), of which most are aborted during the critical phase. Abortion of florets from 6 to 20 days pre-anthesis was strongest in basal spikelets, where on average 91% of the initiated florets are aborted. Contrary to this, in the two central spikelets analysed only 49% and 52% of the florets were aborted during the same time (6-20 days pre-anthesis; Supplemental Table S4). To further understand rudimentary basal spikelet formation, we therefore investigated what causes the disproportionality high abortion of florets in basal spikelets.

### Lower resource availability is an unlikely cause for low grain set in basal spikelets

Previously, Gonzalez (2011) and Stockman (1983) suggested that the increased abortion of florets in the base is due to reduced resource assimilation. This hypothesis is based on the general finding that floret abortion is increased by reducing overall assimilate availability and that basal spikelets have less dry matter weight at anthesis. Furthermore, this hypothesis is supported by the repeated finding that basal spikelets are most reactive to changes in the environment, i.e., loosing or gaining over-proportionately more florets when source strength is altered (Stockman et al., 1983; Savin and Slafer, 1991). Stockman et al. (1983) furthermore suggested that the reduced dry matter weight of basal spikelets indicates their lower assimilate priority. However, as dry matter is measured post-abortion, it cannot be determined if the reduced dry matter weight is due to less resource availability or due to the florets being aborted and thus less tissue growth being supported after their abortion. Stockman et al (1983) are, to our knowledge, also the only study that directly measured carbohydrate concentrations in the apical, central and basal sections of the spike (albeit in controlled environments and with single stemmed plants). Confirming results from their previous study on the whole spike (Fischer and Stockman, 1980), they found that soluble carbohydrate concentrations peak in the spike 12 days pre-anthesis in central and apical spikelets (Stockman et al., 1983). Interestingly, carbohydrates peaked 3 days later in the basal (2^nd^ and 3^rd^) spikelets. They concluded that florets in basal spikelets are more readily aborted as they are, “the sink of lowest priority in the spike” (Stockman et al., 1983). However, this interpretation ignores the fact that basal spikelets accumulate the highest percentage maximum carbohydrate concentration of 30-40% (of the dry matter weight) at 15 days pre-anthesis, greater than the central and apical spikelets which have maximum carbohydrate concentration of 20-25%. At 12 days pre-anthesis, both basal and central spikelets have an equivalent concentration of ~20-25%. Our data supports this re-analysis of their results as we didn’t detect significant differences in sugar concentrations between the central or basal spikelet or rachis, although we also found that there is a tendency of slightly higher sugar concentrations at the base and lowest concentration in the apical region (Table 1, Supplemental Table S5). In conclusion, re-analysis of the Stockman et al., (1983) data and our results do not support the hypothesis that basal spikelets have less resource availability.

### Differences in grain set across the spike can be predicted using floret development stages pre-abortion

To understand the causes of increased floret abortion in basal spikelets we divided the question into two parts. One part concerns the factors that initiate floret abortion and determine the overall degree/strength of abortion. As discussed above, the extent of abortion is largely decided by the availability of resources. The second question concerns the order of floret abortion across the spike, which has been shown to always start in the most distal florets of the spikelet and, if resources are limited, moves inwards within a spikelet. Factors proposed to make distal florets more likely to abort include the distance from the rachis (Kadkol and Halloran, 1988), their size, or the developmental age of the floret at the time of abortion (Ferrante et al., 2020). As the most distal florets of the spikelet are also the youngest and smallest, these factors cannot easily be disentangled.

In 2020, Ferrante et al. showed that the improved grain set in a modern cultivar stemmed from its faster rate of floret development pre-abortion, which improved the survival rate of the more distal florets, compared to the traditional cultivar. Furthermore, they found that lower nitrogen levels reduced floret development rates and thus negatively affected floret survival. The response of the cultivars was linear, meaning that the improved floret development in the modern cultivar was still beneficial under lower nitrogen levels. The study thus connected pre-abortion development of florets to their likelihood of surviving abortion and showed that environmental factors negatively affecting development pre-abortion reduced the survival chance of distal florets (Ferrante et al., 2020). A similar observation had been previously made in barley, where the chance to survive abortion was highly dependent on the development stage attained at the start of floret primordia mortality (Arisnabarreta and Miralles, 2006).

We tested this hypothesis with our data and investigated if reduced development pre-abortion in basal spikelets could be the cause of their increased abortion. We calculated the survival probability of a floret based on its Waddington stage at maximum floret development pre-abortion (20 days pre-anthesis in our 2022 trial, Figure 2). Therefore, the survival probability of a floret was independent of its position along the spike or spikelet. We used the sum of the survival probabilities of the first eight florets within a spikelet as prediction of the number of grains per spikelet. We found that there was a good fit between predicted and actual grain numbers per spikelet if survival probability increased once florets had passed Waddington stage 5.5 (Figure 4). The predictions based on Waddington stage 5.5 were able to capture the gradient of grains per spikelet from the centre to the base of the spike and predicted lower grain values in the basal spikelets compared to central for all genotypes (Figure 4). This aligns with the result of Ferrante et al. (2020) that development stage of florets at abortion is highly relevant for their likelihood of survival and supports our hypothesis that nearly all florets in basal spikelets are aborted due to their reduced development and not due to reduced resource availabilities.

In our study, the closer a floret was to reaching Waddington stage 5.5, the higher its survival chance became. Waddington stage 5.5 was chosen as the previous naïve analysis of all floret data combined suggested that this stage was a good threshold to match grains per spikelet (Figure 3A). Furthermore, we also performed an unconstraint optimisation for each genotype to find the best parameters for the floret survival probability. This allowed us to find the Waddington stage for each genotype that results in the best predictions. Interestingly, these optimised values ranged between 5.0 to 5.7 for the six different genotypes and two locations, highlighting that Waddington stage 5.5 is indeed an important stage to be reached for high floret survival.

Reanalysis of the Ferrante et al., (2020) data also lends support to Waddington stage 5.5 as the onset of floret abortion and important checkpoint. At this stage (equivalent to −270 °C d from anthesis in their study), all of the florets that reached 5.5 completed development up to anthesis, and conversely 91% of the florets that did not reach anthesis also had not reached 5.5 at −270 °C d (Supplemental Figure S2). For the different genotypes and conditions investigated in our study and in Ferrante et al., (2022) Waddington stage 5.5 gave good predictions, suggesting that it is rather the developmental rate pre-abortion that changes between genotypes and conditions, rather than the Waddington stage checkpoint itself. Further studies would be warranted to elucidate the overall applicability of this hypothesis. Investigations of the molecular mechanisms happening when a floret reaches Waddington stage 5.5 will be important next steps to understand the importance of this stage for floret abortion.

The initial and optimised grain predictions were both able to predict the gradient across the spike, however some spikelets along the spike were predicted worse than others. Focusing on the predicted values using the common mean value of 5.5, the grains in central spikelets of some genotypes were under-predicted. Using the optimised predictions, grains per central spikelet were predicted highly accurately, however the predictions for the 1^st^ and 2^nd^ most basal spikelet were significantly different from the actual grain numbers (Figure 4). This suggests that even though the predictions based on Waddington stage pre-abortion can account for the majority of the observed variation, additional factors likely play a role in determining abortion differences in the central versus the basal spikelet.

Despite these potential shortcomings, using this very simple rule we were able to predict the grains per spikelet in basal and central spikelets using a general framework. This suggests that the signalling pathways of floret abortion might be the same in central and basal spikelets. The severe delay in floret development in basal spikelets from spikelet initiation until stem elongation might thus explain rudimentary basal spikelet formation, rather than the previously proposed hypothesis of reduced resource availability in basal spikelets. A similar mechanism has been proposed in barley. Unlike wheat, barley has determinate spikelets and an indeterminate spike and therefore spikelet abortion has been studied in much more detail (Alqudah and Schnurbusch, 2013). Under salt stress conditions, all growth stages pre-abortion are shortened and spikelet growth and development is diminished, leading to increased abortion of apical and basal spikelets (Boussora et al., 2019).

### Improvement of basal spikelet fertility through targeting pre-abortion development

It has been proposed that a reduction in the variation in spikelet fertility across the spike could be a promising avenue to increase yields and improve grain size homogeneity in breeding programs of small cereals grains (Arisnabarreta and Miralles, 2006; Philipp et al., 2018). Our results suggest that this would not be possible by reducing abortion, but rather through improving spikelet and floret development pre-abortion. Reducing the amount of abortion either through improved genetics (such as *GNI1* introgression; Sakuma et al., (2019)) or increased resource availability leads to increase grains per spikelet. However, this seems to always be in a linear fashion across the spike and would not specifically improve basal spikelets compare to central.

Interestingly, when the survival probability function was left to vary freely during optimisation, the resulting means of the survival probability function all fell within a range of 5.0 to 5.7, consistent with our hypothesis that Waddington stage 5.5 is an important developmental stage for floret abortion survival. Overall, the differences in mean values found by optimising the function for each genotype matched the flowering dates of the varieties. The CDF mean Waddington stage value was highest in Skyfall and Cadenza (in CF trial), followed by Robigus, Ruskin, *VRT-A2a* and *VRT-A2b* (Figure 4D), which matches the sequence of flowering dates of these genotypes scored in control plots in 2022 CF trials (Supplemental Table S1), with Skyfall flowering first and *VRT-A2b* flowering last. This suggests that Skyfall florets were more advanced and thus the common mean of 5.5 overpredicted grains. Skyfall also appeared to be slightly ahead of the other genotypes in floret development as it was already at maximum floret potential a week before the other genotypes (Figure 2). Genetically mapping the effect of Skyfall on RBS would thus potentially be able to uncover further genes involved in the control of pre-abortion development.

In this study, we included a set of NILs for *VRT-A2*, a MADS-box transcription factor, previously shown to increase RBS numbers under controlled and field conditions (Backhaus et al., 2022). We confirmed that RBS was increased by *VRT-A2b* and furthermore found the allele to have no interaction with shading (Figure 1). Using the same parameters, we were able to predict the grains per spikelet as accurately as for the wildtype *VRT-A2a* NIL. This suggests that the introgression of *VRT-A2b* indeed affects pre-abortion development rather than increasing abortion *per se* in the basal spikelets. This is also supported by the subtle overall delay in development across florets in *VRT-A2b* NILs by 0.21 Waddington stages compared to the wildtype at the onset of abortion (Supplemental Table S8). Optimisation of the cumulative distribution function furthermore came to very similar mean values for both NILs in CF (*VRT-A2a* = 5.08 and *VRT-A2b* = 5.04) as well as in Morley (*VRT-A2a* = 5.65 and *VRT-A2b* = 5.44).

In Morley, we only collected data for the NILs and the optimisation found a better fit for both NILs using a more advanced Waddington stage mean than in CF (Figure 1E). This might suggest that the plants were marginally more advanced in development in Morley, although flowering dates are not available for this experiment to explore this hypothesis. The differences in mean Waddington stages to accomplish better fits for individual genotypes might be mainly correcting for differences that arose because we sampled all genotypes on the same day. Thus, we did not account for developmental differences between the genotypes as there may have been varietal variation, as suggested by the differences in flowering dates (Supplemental Table S2).

Our study highlights the importance of the ‘critical phase’ of wheat development, adding basal spikelet abortion to the traits affected during this phase. Reducing abortion during this phase is a promising avenue for future yield increases and as the spike is particularly sensitive to resource limitation in the 10-20 days preanthesis. Management practices might be a promising tool to reduce abortion as precise application of fertiliser at maximum floret stage could reduce abortion. Ferrante et al. (2020) showed that reduced nitrogen application throughout the growth season (by 75%) slows down the development of florets, which leads to an increase in florets that are not advanced enough to survive abortion. The same effect of reduced nitrogen on floret survival had previously been proposed by Abbate et al. (1995), however they only recorded the effect of nitrogen reduction on grains/m^2^ and did not dissect the trait further. It would be interesting to investigate if applying important signalling compounds of resource availability, such as sucrose or trehalose 6-Phosphate (Paul et al., 2018), during the time of floret abortion could reduce abortion rates and thus counter-act increased floret abortion in low nitrogen conditions.

In this study we found that it is the reduced development of floret primordia pre-abortion in basal spikelets that can explain the increased loss of florets during abortion. In several genotypes, all florets are aborted in the basal spikelet, which we propose to be the reason for their rudimentary appearance in the mature spike. Thus, it would be the initial establishment of the basal spikelets and their development rates pre-abortion that need to be targeted to improve homogeneity across the spike.

## Materials and Methods

### Genetic material and plant growth

Wheat germplasm used in this study includes hexaploid UK wheat cultivars Cadenza, Robigus, Ruskin, Skyfall, and near isogenic lines (NILs) differing for the *P1* locus described in Adamski et al. (2021). We used two sibling BC6 NILs with Paragon as the genetic background, differing for the presence of the wildtype *VRT-A2a* allele or the *T. polonicum VRT-A2b* allele. We evaluated cultivars and NILs in three field experiments. One trial was located at The Morley Agricultural Foundation trials site, Morley St Botolph (M), UK (52°33’15.1”N 1°01’59.2”E) in 2021/22. Two trials were located at the John Innes Centre Experimental trials site in Bawburgh (Church Farm; CF), UK (52°37’50.7”N 1°10’39.7”E) sown in 2020/21 and 2021/22 in two different field locations. We drilled all experiments as 1.2 m^2^ plots (1 m x 1.2 m) and we sowed them by grain number for comparable plant densities aiming for 275 seeds*m^2^. We treated all trials with herbicides and fungicides as needed and we applied between 211-218 kg of nitrogen per hectare and 72-75 kg of sulphur per hectare over the growth season. In the 2021/2022 season at Bawburgh (Church Farm; CF) we only applied 50kg sulphur per hectare. The experiment was also irrigated once, on the 27^th^ of April 2022, at a rate of 12mm per hectare, as the season and field were extremely dry. We arranged plots in a randomized complete block design (RCBD) with a split plot arrangement (main plot = Shading treatment, sub plot= Genotype) and three replications.

### Shading treatments

We applied shading by covering the main plots (6 genotypes) with 55% Shade Netting made from long-lasting HDPE tape monofilament threads (LBS Horticulture, product ref: NETS001). Nets were cut to length and supported by metal cones and a bamboo frame, installed ca 20 cm above the crop canopy, while avoiding the netting to touch or constrain stem/spike growth. We pulled nets down and secured on the sides (with reusable zip ties) to reduce light entering from the sides. We measured the relative light penetration of the net using a light meter (Skye Instruments Ltd SKP-200; Supplemental Table S1). In 2021 (CF), we applied two shading treatments for 12 days each. Shading A was applied from 16.05.2021 to 28.05.2021, whereas Shading B was applied from 29.05.2021 to 10.06.2021. We chose dates based on estimation of anthesis happening circa 2 weeks after shading applications. In 2022, we applied three shading treatments and overlapped each other by one week. Shading C was installed on 05.05.2022 in CF2022 and 06.05.2022 in Morley (M2022). Shading D was installed on 12.05.2022 and 13.05.2022 in CF2022 and M2022, respectively. Shading E was installed on 19.05.2022 in CF2022 and M2022. We removed all shading treatments 13 days after installation, except shading D in M2022, which was removed after 12 days.

### Phenotyping

To assess floret development during the 2022 season, we cut one main tiller (below the spike, which can be easily assessed by finding the last internode) from the central area of the plot and placed in an emptied box of 1000 μl tips filled with water. We marked the remaining plant to avoid sampling from the same or neighbouring plants again. For each genotype, we took one spike from each block in the control and shading treatment as described in Figure S3. For each spike, we dissected the six most basal and two central (most developed) spikelets, and we determined the number of living floret primordia per spikelet and the Waddington stage of each floret, using the Waddington scale and images from Prieto et al. (2018) as reference.

We collected samples at each of the five timepoints depicted in Figure S3 and coinciding with the installation/removal of a shading treatment. For the first timepoint (27 days pre anthesis) we collected only spikes from the control plots the same day as Shading C was applied and all subsequent collection timepoints were in one-week intervals. We collected samples each week from the control plots. For Shading C and D, we collected samples one and two weeks after start of the treatment. Samples were only collected after two weeks of Shading E (Figure S3).

At the end of the growing season, we hand-harvested mature plants. In Morley we collected grab samples of 10 main spikes from each plot as the ground was too hard to pull plants. For each plot in CF (2021&2022) trials, 10-20 individual plants were pulled from the centre of the 1.2 m^2^ plot, which allows for more accurate separation of main and side tillers. Plants were separated, and roots were removed 5 cm above the crown. We assessed plant dry weight before removing all spikes in 2022 but after removing spikes in 2021. We also recorded spikelet number and the number of grains for each spikelet across the main spike. For Morley (2022) we processed spikes in the same manner as in CF trials.

### Sugar measurements

Sugar samples were collected by sampling three spikes per plot in the corresponding timepoint. For CF 2021, we sampled at the end of shading B, whereas for CF 2022 and Morley 2022, we collected spikes at the end of shading D from control and shading plots. For each replicate, we dissected spikes into four basal spikelets and four central spikelet and furthermore separated the rachis of these. Apical spikelet and rachis were collected only in 2022 field experiments. Immediately after dissection the tissues were snap frozen in liquid nitrogen. Samples were consequently stored at −80C until further processing. Samples were ground using pestle and mortar and ~20 mg of powder (exact weight was recorded) was dissolved in 1.2 mL of ethanol 80% (v/v) in screw-capped tubes. These extracts were mixed thoroughly and incubated for 1 h at 80 °C, mixing halfway through. Extracts were subsequently centrifuged at 12000xg for 1 min and supernatant was collected. We stored samples at −20 °C until assayed.

The method of sugar extraction and measurement is based on Griffiths et al. (2020). Briefly, to perform the assay, we added 5 μL of ethanolic extract to 145 μL of reaction buffer [100 mM HEPES pH 7.4, mM MgCl2, 1 mM NAD+, 0.5 mM ATP, 1.5 U/μL hexokinase]. We first measured baseline absorbance at 340 nm. Subsequently, different enzymes are added sequentially for measurement of either glucose or fructose. The first reaction is initiated by addition of 1.2 U of glucose-6-phosphate dehydrogenase from *Leuconostoc mesenteroides* and incubated for 45 min. Then, absorbance at 340 nm is measured to determine glucose concentration. Subsequently, 0.2 U of phosphoglucoisomerase from yeast are added and the reaction is carried out for 46 min at room temperature, to determine fructose concentration. Finally, 10 U invertase are added and reactions incubated for 1.5 h, to determine concentration of sucrose. All reactions were performed at room temperature in flat-bottomed 96-well microtiter plates and the measurement of each sugar was done by measuring the reduction of NAD+ to NADH at 340 nm (Varioskan LUX, ThermoFisher for 2021 trials; Spectramax, Molecular Devices for 2022). To correct for variation between runs, a calibration curve for the three sugars was included in each plate and the concentrations were calculated by interpolation. Reactions were performed in triplicates and normalised to the weight of the powder used in extractions. In 2021, all sugar measurements were performed at JIC. In 2022, all measurements were performed at Rothamsted Research Institute.

### Data processing and analysis

Using the raw phenotypic data from mature spikes, we calculated the number of rudimentary basal spikelets (RBS), total spikelets, and central spikelet fertility in R. We defined RBS as spikelets carrying no grain and we determined RBS for each spike individually. On average, spikes had ~20-25 spikelets, we therefore calculated the number of grains per central spikelet by averaging the number of grains in the 10-ear samples from the 10^th^, 11^th^, and 12^th^ spikelet (from the base). Using the raw floret development scores, we calculated the total number of florets per spikelet by counting the floret Waddington scores per spikelet.

To determine the differences between the genotype and treatments, we performed analysis of variance (ANOVA) on mature plant and floret development phenotypic data. For the analysis of mature plant data from individual field experiments, we used a split-plot two-way ANOVA including genotype as sub-plot and shading treatment as primary plot (performed in R RStudio 2022.02.0, using ‘agricolae’ package (version 1.3-5; De Mendiburu Delgado (2009)) sp.plot() function and for post-hoc multiple comparisons the LSD.test()). Floret survival was analysed using R base ANOVA function and post-hoc Sidak test. To analyse differences in sugar concentrations across the spike sections (Apical, Central and Basal), we first performed a 3-way ANOVA in R (RStudio 2022.02.0, using ‘agricolae’ package (version 1.3-5; De Mendiburu Delgado (2009)). If significant interactions were detected the effect of ‘section’ was analysed by sub-setting the data by ‘tissue’ or ‘treatment’ (depending on which had significant interactions with ‘section’). Each field experiment was analysed individually. ANOVA and post-hoc tests performed are indicated below each supplemental table. Convidence interavals (CI) were calculated in R by determining the mean (x) estimate and subtracting (LCI) and adding (UCI) the variation in the estimate:

To predict the survival chances of each floret, we utilised the cumulative distribution function (pnorm(), R RStudio 2022.02.0). Initially, we calculated survival probabilities using a fixed mean of 5.5 and standard deviation (SD) of 0.195. We applied this function to florets 1 to 8 for spikes collected 20 days pre-anthesis (i.e., maximum floret stage) in control conditions in CF (all genotypes) and Morley (only NILs). The probabilities of florets 1 to 8 within each spikelet were summed to get the predicted number of grains per spikelet. The idea being that if the survival chance on all florets is very high (nearly 1), the number of grains would equal number of florets per spikelet. We used 5.5 as a mean value for the cumulative distribution function as our hypothesis was that Waddington 5.5 is an important stage, florets beyond this stage thus have a very high survival chance while florets below have very low survival probability. We chose SD = 0.195 as this was the standard deviation of the grains/spikelet dataset from mature spikes.

We used the optim() function in R, with the choice of optimisers defined by method=“SANN” (simulated annealing) and method=“Nelder-Mead” (downhill simplex or Nelder-Mead method), with an objective function (with parameters mean and SD of the cumulative normal distribution) that defines the Euclidean distance between the predicted and measured grain number. The CDF mean and SD were optimised to minimise the difference between predicted and measured grain numbers.

## Acknowledgments

We thank the JIC Field Experimentation team, Tobin Florio, Sophie Eade and Pamela Crane for technical support in field experiments. We thank Desislava Kostadinova for her support during the 2021 field sample collection.

## Author Contribution

(Contributions are based on the CRediT Contributor Roles Taxonomy)

AEB: Conceptualization, Methodology, Data curation, Formal analysis, Investigation, Visualization, Project administration, Supervision, Writing - original draft

CG: Investigation, Methodology, Formal analysis

AVC: Formal analysis, Investigation

JS: Methodology, Resources

RL: Methodology

RM: Software, Formal analysis

CU: Conceptualization, Funding acquisition, Methodology, Project administration, Supervision, Writing - review & editing

## Supplemental Table and Dataset Legends

**Supplemental Table S1:** Solar radiation (umol/sec/m2) measured in CF 2021 control and shading plots under various light conditions.

**Supplemental Table S2:** Flowering dates of all genotypes analysed in this study in Church Farm (CF) trials in 2021 and 2022.

**Supplemental Table S3:** Statistical analysis of phenotypic data, performed individually for each field experiment.

**Supplemental Table S4:** Number of florets pre-abortion (20 DPA) and the number of florets aborted from 20 DPA to 6 DPA abortion in control, and in control versus Shading D plots.

**Supplemental Table S5:** Statistical analysis of sugar concentrations (μg/mg tissue weight) in dissected spike tissues, performed individually for each field experiment.

**Supplemental Table S6:** Average development stage (Waddington) of floret 1-8 in basal (B) six and central (C) two spikelets across all genotypes 20 DPA (Control treatment only).

**Supplemental Table S7:** T-test comparison of predicted versus actual grain values per spikelet

**Supplemental Table S8:** Average development stage (Waddington) of florets 1-8 in basal (B) six and central (C) two spikelets 20 DPA (Control treatment only) for *VRT-A2* NILs.

**Supplemental Dataset S1:** All raw field data collected in 2021 and 2022 for mature spikes (post-harvest).

**Supplemental Dataset S2:** All developmental scores taken for floret development in the basal six and central two spikelets from spike collected in CF 2022.

**Supplemental Dataset S3:** Raw normalized sugar concentration (μg/mg tissue wt) for samples collected in CF 2021 and 2022.

**Supplemental Dataset S4:** Number of grains predicted for each spikelet using CDF, with and without optimisation.

## Supplemental Figures

**Supplemental Figure S1:**
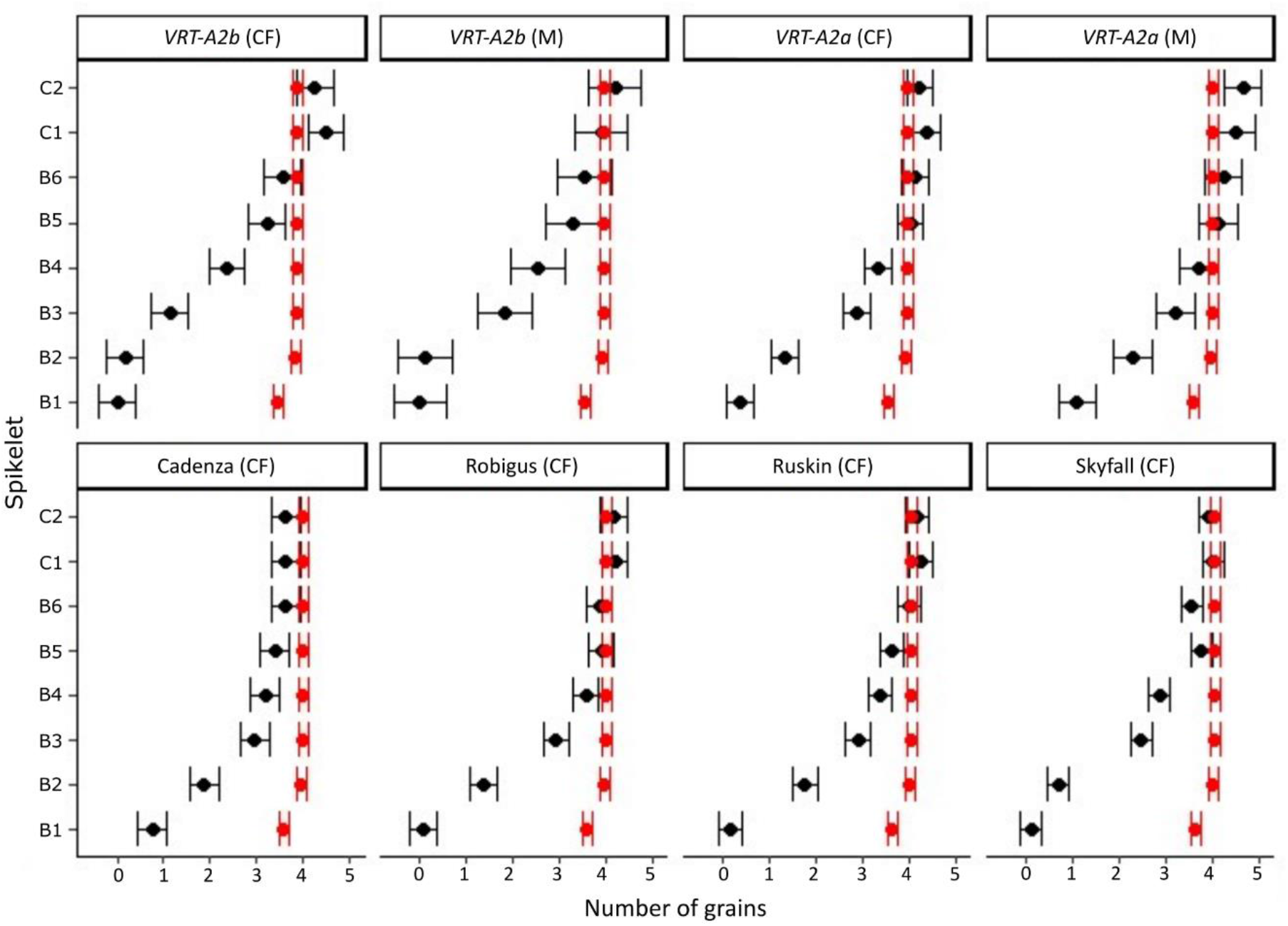
Predictions of grains per spikelet using floret count per spikelet pre-abortion (20 DPA) and deducting same number of florets per spikelet (4 florets) across all spike positions. Black = Number of grains per spikelet recorded in mature spike, Red= predicted grains/spikelet. B = Basal (spikelet), C = Central (Spikelet).

**Supplemental Figure S2:**
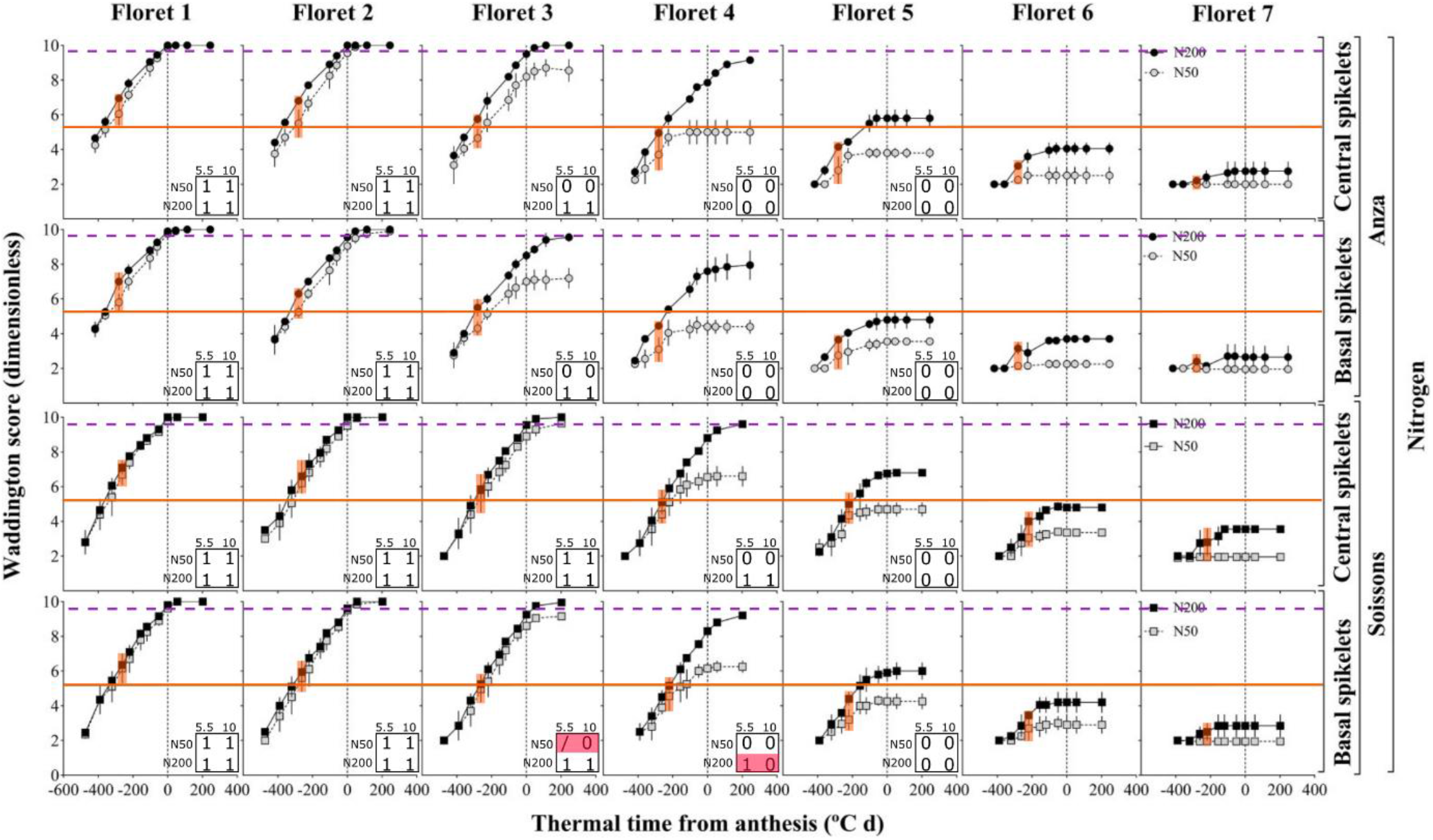
Re-analysis of Ferrante et al (2020) Figure 9. Orange bar was added to mark Waddington stage 5.5 and the data points corresponding to maximum floret number stage pre-abortion (circa −270 °C d) were highlighted in orange to determine if florets had reached Waddington stage 5.5 at that time. Purple dotted line indicates cut-off for florets considered to have reached maturity (W10). Out of the 56 floret development traces, 22 reached Waddington stage 10 by the end of the time course and all of these had passed Waddington stage 5.5 at −270 °C d (which corresponds to maximum floret number stage according to Figure 8 (Ferrante et al., 2020)). Of the 34 florets that did not reach Waddington stage 10 at maturity, 91% had not reached W5.5 at −270 C°. Only floret 4 of the basal spikelet in Soisson (high nitrogen) had reached W5.5 but then did not develop until the W10 cut-off (purple line) while the development stage of floret 3 in the same genotype and spikelet at nitrogen level 50 could not be accurately scored. Decisions on floret development are indicated in box on bottom right of each panel. If florets reached the stage they were scored 1, if they didn’t they were scored 0. Red shading indicates that floret development at W5.5 does not match their development stage at W10. As all florets were infertile beyond floret 5 we stopped analysis beyond floret 5. Original legend from Ferrante et al (2020), *“Dynamics of the floret development from floret 1 (F1, floret primordium closest to the rachis) to floret 7 (F7, floret primordium most distal to the rachis) in each of the two spikelet categories considered of the main-shoot through thermal time from anthesis (negative values represent the period before anthesis) in the N experiments for Anza and Soissons. Grey and black symbols correspond to N50 (50 KgN ha^-1^) and N200 (200 KgN ha^-1^). Each data-point is the average of all replicates across two growing seasons and within each replicate the value was the average of 10 (2010-11) and 5 plants (2011-12), bars represent the standard error of the means (not visible in some cases as it was smaller than the body of the symbol).”*

**Supplemental Figure S3:**
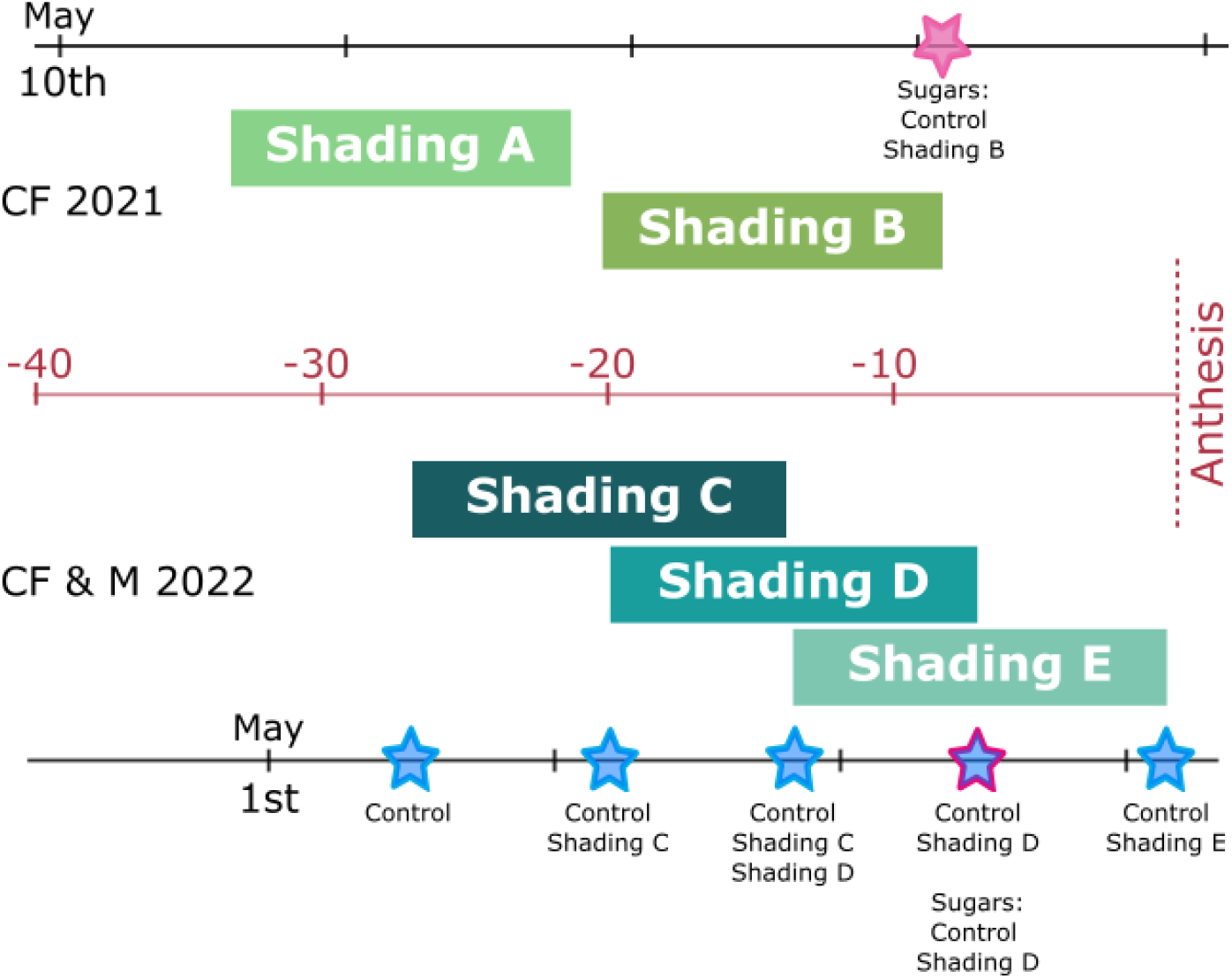
Schematic of floret development and sugar sampling in 2021 and 2022. Blue stars indicate floret collection dates, pink star or outline indicate sugar collection dates.

